# Genome-scale mRNA transcriptomic insights into the mature growth dynamic of Cigar tobacco leaves

**DOI:** 10.1101/2022.09.26.509548

**Authors:** Jinpeng Yang, Di Wan, Baoming Qiao, Jun Yu, Zongping Li, Daisong Liu, Peijun Lv, Jinwen Hu, Xiongfei Rao, Fangsen Xu, Sheliang Wang, Chunlei Yang

## Abstract

The harvest time is a key factor for cigar leaves with high quality, which varies greatly depending on the environment. Here, we performed a genome-scale mRNA transcriptomic analysis on the cigar cultivar CX-26 (Nicotiana tabacum L.) to evaluate the relationship between gene expression and growth state. The leaves were harvested with 67 (T1), 71(T2) and 75 (T3)-day growth. A total of 80,502 genes were detected in the CX-26 leaves, of which 64,611 genes were annotated on the reference genome. Principal component analysis showed that T1 and T2 leaves had a high overlapping pattern, while T3 leaves were distinct. Indeed, T1 and T2 leaves had fewer differential expressing genes (DEGs), while T3 leaves had 26,456 DGEs from T2 leaves, supporting the distinct growth of T3 leaves. GO annotations mainly enriched the photosynthesis-related metabolic process, catalytic activity and binding biological processes. KEEG analysis identified the key pathways including photosynthesisantenna proteins, plant hormone/MAPK signaling pathway and plant-pathogen interaction. The maturity regulation and defense response-associated hormones abscisic acid and jasmonate acid were higher in the T3 leaves than that in T1 and T2 leaves confirming the KEEG analysis. Furthermore, several photosynthesis-related enzymes and a transcription factor were highlighted in the gene regulatory network, which might regulate the dynamics of carbohydrate metabolism, lipid metabolism and energy metabolism. In summary, our study provides insight into the growth state of CX-26 cigar leaves.

## Introduction

Plant leaf originates from the leaf primordia in the shoot apical meristem (SAM) site. The mature leaf morphology results from the development transition of the young leaf that undergoes leaf primordia growth along with the cell proliferation and polarity to a maximal size [1]. A mature leaf has strong photosynthetic activity upon the function of chlorophylls and carotenoids. Senescence is the last phase of leaf growth (post-mature leaf) and plays an important role in plant growth and development such as the reuse of mineral nutrients [2]. The senescence was characterized by the degradation of chlorophylls with a series of enzyme catalytic reactions [3].

Maturity is pivotal for the quality of tobacco leaves depending on the accurate judgment of harvesting time [4]. To explore a fast quantitative method on the maturity of flue-cured tobacco leaves, researchers established the mathematical relationship between the leaf HSV (Hue Saturation Value) color value and chlorophyll content, and chlorophyll content and SPAD value, respectively [5]. Based on this, a relation model TMDHSV was set up to effectively evaluate the leaf maturity. Recently, an improved lightweight network architecture for identifying tobacco leaf maturity was reported, which is based on the deep learning of artificial intelligence [6]. Based on these tools, the quality of flue-cured tobacco leaves was greatly improved. Changes in the appearance of the leaf are the consequences of physiological processes. A time-series transcriptomic analysis combining metabolomic profiling in flue-cured (K326 variety) tobacco leaves identified important regulatory networks involved in photorespiration and the tricarboxylic cycle, therefore might regulate the C/N balance during leaf development [7].

It has been reported that either premature leaf or post-mature leaf cannot possess appropriate coordination of chemical contents and aroma components in the cigar, leading to a reduction in the quality of cigar leaves [8]. However, the maturity judgment of cigar leaf usually depends on the experiences of growers. In particular, the literature on regulatory networks between maturity and gene expression in cigar leaf in terms of genome-scale remains elusive. Here, we performed a genome-scale transcriptomic analysis of the middle leaves of a cigar wrapper variety CHUXUE-26 (CX-26) harvested at different times. Transcriptomic results point to the important regulatory networks associated with the mature growth state of CX-26 leaves.

## Results

### Dynamics of chlorophyll and moisture content in CX-26 leaves

To estimate the growth state changes of cigar leaves, we measured the moisture contents and SPAD values of cigar leaves at three time points (July 22 (T1), 26 (T2) and 30 (T3)), firstly. The moisture contents showed a gradually decreasing trend (Fig 1A). However, the isolate values of three measurements were very close. To observe the dynamics of chlorophyll contents at three time points, we recorded the SPAD values (Fig 1B). Similarly, the older leaves had lower SPAD values among them, and the reduced trend was greater than that of the moisture content. These results suggest that T3 growth state of CX-26 cigar leaves was largely changed away from T1 and T3 leaves.

**Fig 1.**
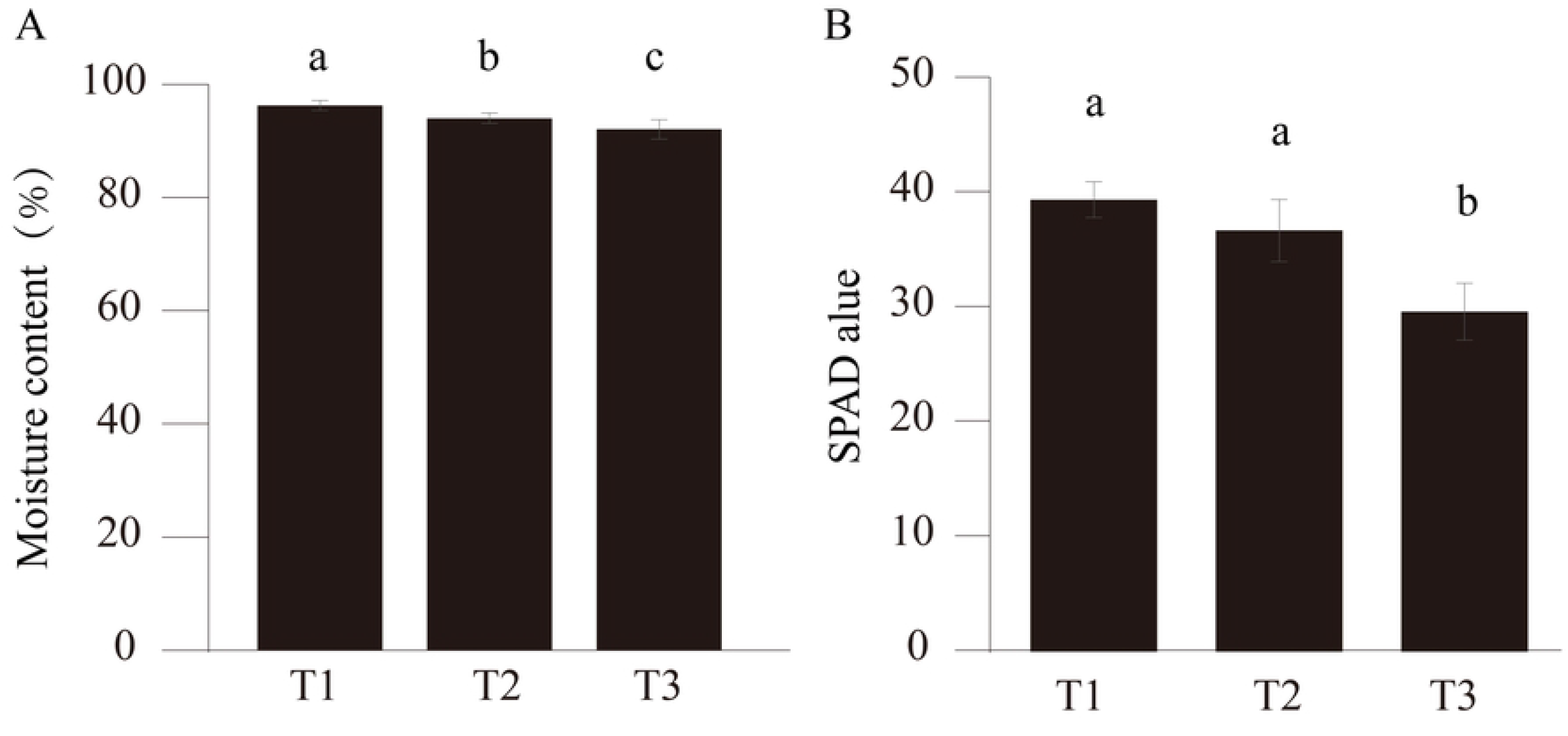
Moisture contents and SPAD values of CX-26 leaves. A, Moisture content analysis of CX-26 leaves. Fresh weights and dry weights of T1, T2 and T3 harvested at different times were used for Moisture content calculation. n=4. B, SPAD values analysis of T1, T2 and T3 leaves. n=4. The significant differences were analyzed with one-way ANOVA.

### General transcriptional changes and differential expression genes in CX-26 leaves

The CX-26 leaves harvested at three time points were used to be RNA-seq analyzed. Four biological samples for each time point (T1, T2 and T3) were selected. A total of 64,611 annotated genes were detected with expression levels, accounting for 80.26% of annotated genes in all databases (Fig 2A). Besides, a total of 15,891 genes without annotation were also detected. All genes detected cover 79.37% of the reference genes. Both genes and transcripts had consistent expression distribution among three groups (Fig 2B-C). The numbers of co-expression genes among T1/T2, T2/T3 and T1/T3 leaves were 45,933, 38,006 and 38,339, respectively (Fig 3). To evaluate the expression correlation among three groups, we conducted the principal component analysis (PCA) of these genes. Interestingly, T1 and T2 leaves showed an overlapping pattern and T3 leaves had an independent expression pattern (Fig 4A). Consistent with this, T1/T2 comparison identified only 159 differential expression genes (DEGs), while T2 /T3 and T1/T3 comparisons had 26,456 and 23,921 DEGs, respectively (Fig 4B). These results indicate that T1 leaves and T2 leaves had a similar growth state while the growth state of T3 leaves largely differed from T1 and/or T2 leaves.

**Fig 2.**
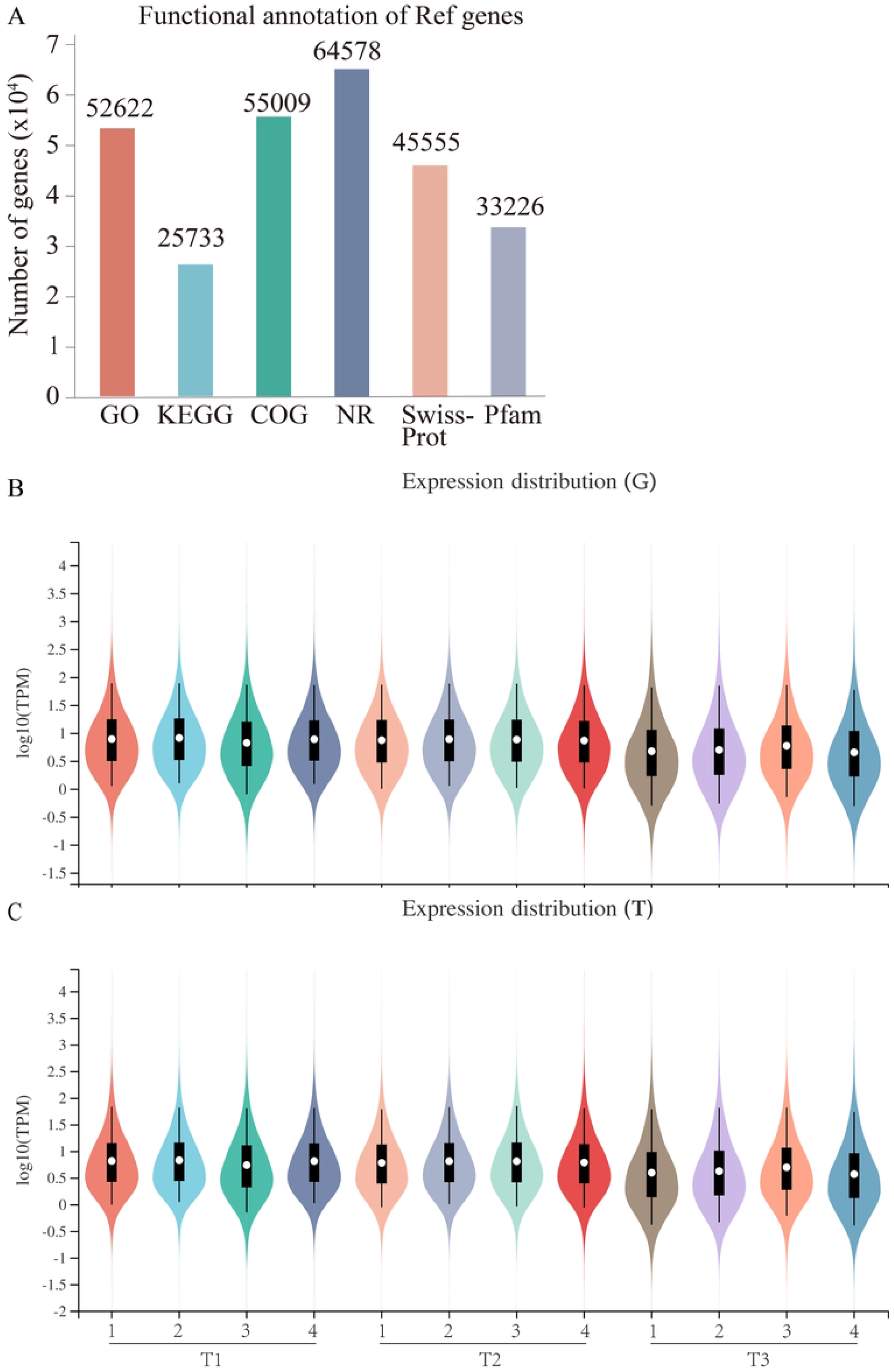
Gene functional annotation and expression distribution of CX-26 transcriptome. A, Numbers of functional annotation genes detected in CX-26 leaves. Clean reads of T1, T2 and T3 transcriptome were aligned to reference genome in six databases (GO, KEGG, COG, NR, SwissProt and Pfam). B and C, The expression distribution of genes and transcripts in CX-26 leaves. n=4.

**Fig 3.**
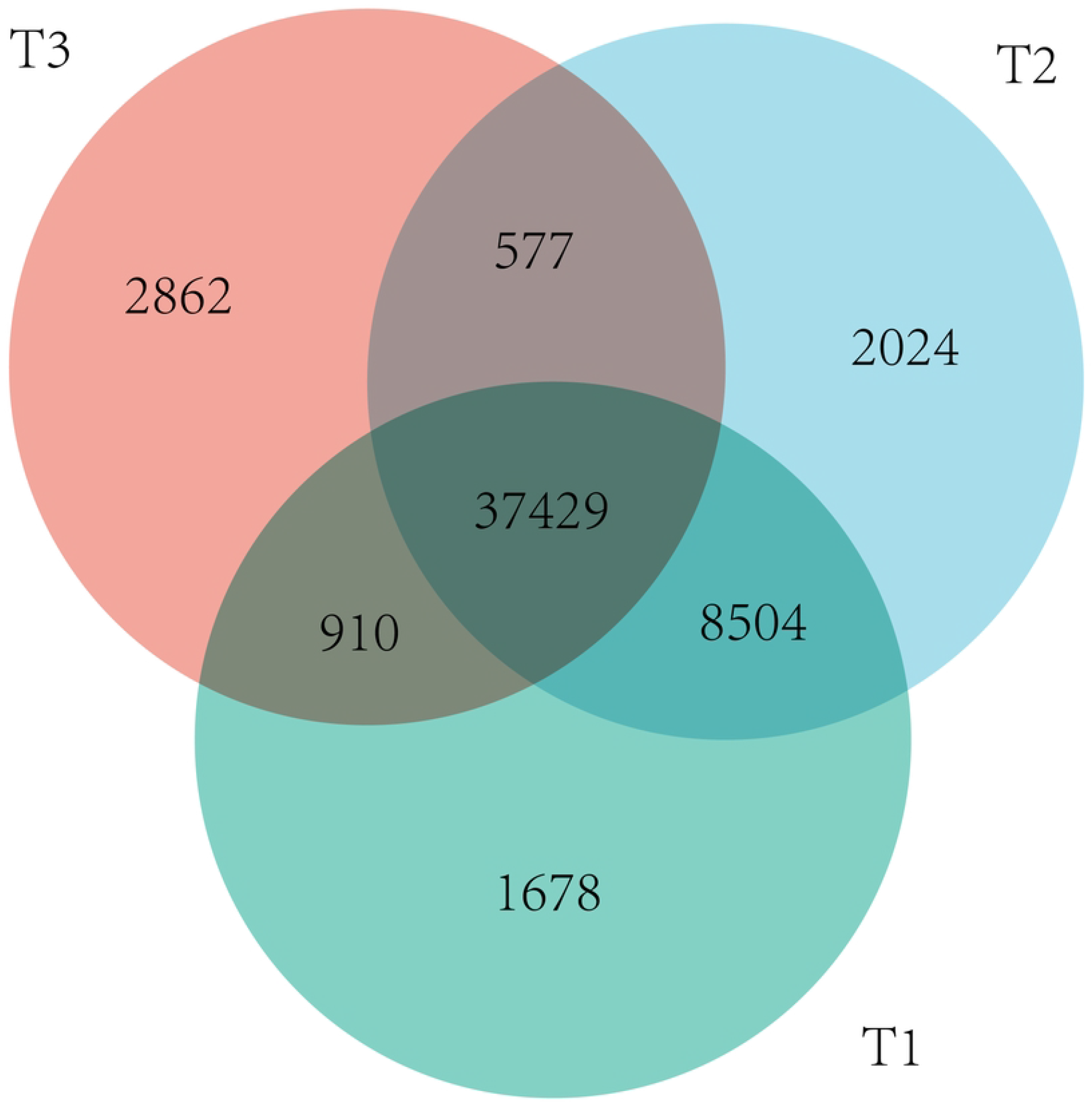
Venn diagram of the numbers of genes detected among T1, T2 and T3 leaves. The numbers of detected specifically in each leaf, co-expressed were indicated in diagram, respectively.

**Fig 4.**
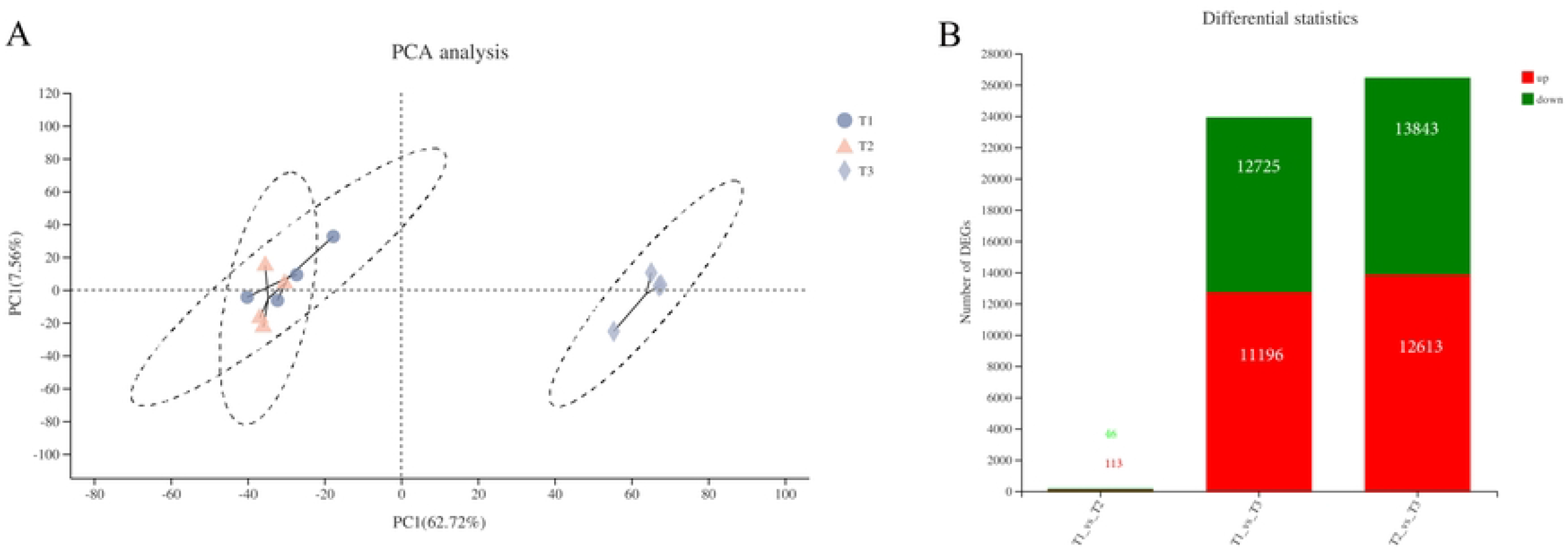
Principal component analysis and differential expression genes in CX-26 leaves. A, Cluster together indicates that T1 and T2 leaves had similar expression profiles, while T3 leaves had a distinct expression profile. B. Number of genes that differentially expressed between each sample was statistically calculated.

### Gene ontology analysis insights into the importance of photosynthesis in mature growth

We further evaluated the growth state-associated molecular function, cellular component and the biological process by gene ontology (GO) annotations analysis among T1, T2 and T3 leaves. Because the T1 and T2 leaves had fewer DEGs, it was hardly annotated in the GO analysis compared with T1/T3 DEGs and T2/T3 DEGs (Fig 5A). In the molecular function, both T1/T3 DEGs and T2/T3 DEGs had strong annotation on the cellular process and metabolic process. In the cellular component, both T1/T3 DEGs and T2/T3 DEGs had strong annotation on the cell part, the membrane part. In the biological process, both T1/T3 DEGs and T2/T3 DEGs had strong annotation on the binding and catalytic activity. This result suggests that the T3 leaves have distinct physiological dynamics from T2 leaves. In details of the T2 and T3 comparison, the top 10 enriched GO items mainly included photosynthesis, light harvesting in photosystem I (59 upregulated DEGs), response to high light intensity (35 up vs 1 down DEGs) regulation of photosynthesis (32 up vs 2 down DEGs), starch biosynthesis process (27 up vs 5 down DEG), photosystem II assembly (24 up vs 2 down DEGs), protein repair (23 up vs 1 down DEGs), fructose 1, 6-bisphosphate metabolic process (20 up vs 2 down DEGs), fructose metabolic process (15 up vs 5 down DEGs), photosynthetic electron transport in photosystem I (19 up DEGs) and pyridoxine metabolic process (11 up vs 1 down DEGs) (Fig 5B). Interestingly, the upregulated DEGs accounted for the main part of each GO item. This result indicates that growth state dynamics from T2 to T3 leaves are tightly associated with the photosynthetic gene regulation process.

**Fig 5.**
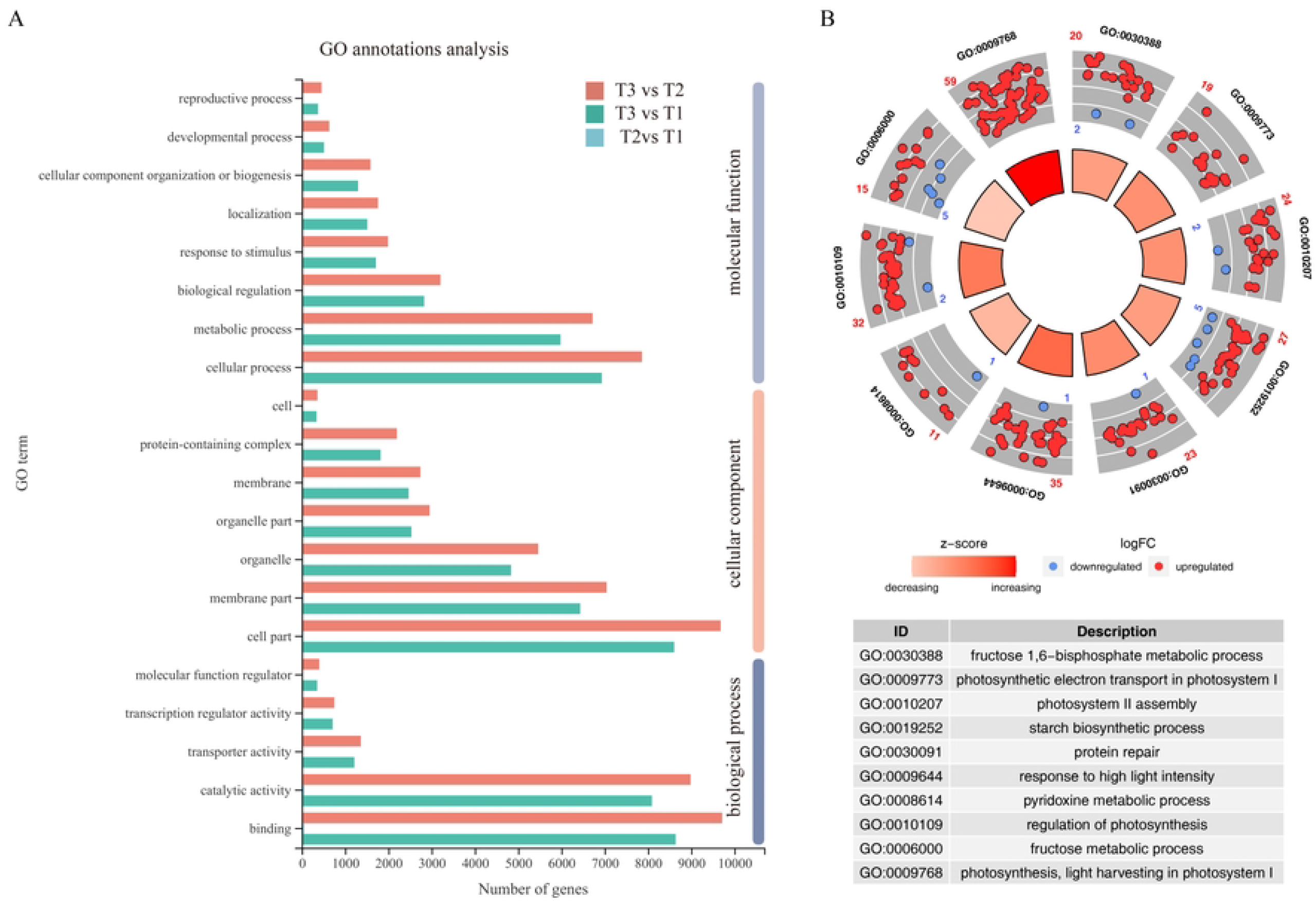
The annotation results of CX-26 leaves transcriptome. GO (Gene Ontology) annotation results of DEGs of CX-26 leaves. The annotated DEGs were classified into molecular function, cellular component and biological process categories (A) and corresponded to GO annotation in T3 leaves (B).

### KEEG analysis and gene regulatory network analysis reveal

#### key pathways involved in growth state dynamics

Furthermore, the KEGG enrichment analysis of T2/T3 DEGs (Fig 6) showed the highest rich factor on the photosynthesis antenna proteins and had the most genes enriched KEEG pathways on plant hormone signal transduction (354 DEGs) followed by plant-pathogen interaction (263 DEGs) and MAPK signaling pathway (243 DEGs) (Fig 6). The gene expression level analysis showed that all the light-harvesting chlorophyll protein complex (LHC) genes had upregulated expression in T3 leaves compared to T2 leaves (Fig 7A). Therefore, the hormonal signaling (FigS 1) and MAPK signaling-mediated environmental biotic or abiotic responses (FigS 2) might be the two key aspects regulating the growth dynamics of T3 leaves including the photosynthetic process and plant-pathogen interaction (FigS 3). It has been demonstrated that abscisic acid (ABA), salicylic acid (SA), and jasmonate acid (JA) positively regulate leaf senescence through activating the expression of Chlorophyll catabolic genes which mediate the chlorophyll degradation [9-11], while IAA inhibits chlorophyll loss [12]. Given the reduction of SPAD value in T3 leaves (Fig 1B), we, therefore, measured the ABA and JA levels in the cigar leaves. T1 and T2 leaves had comparable ABA levels, and T3 leaves had higher ABA levels than that T1 and T2 leaves (Fig 7B). A gradual increase consequence was observed from T1 leaves to T3 leaves (Fig 7C). These results suggest that T3 cigar leaves enhance their hormonal response in the growth dynamics. We further conducted the gene regulatory network to evaluate the gene regulatory relationship underlying mature growth dynamics. The top 7 regulatory gene hubs (Fig 8) are the gene_42749 encoding dehydroascorbate reductase 1, gene_222 (function unknown), gene_1002 (starch synthase 2), gene_53955 (glycogen/starch synthases), gene_75915 (starch synthases), gene_7934 (2-phosphoglycolate phosphatase 1) and gene_77328 (bZIP transcription factor family protein). Except for the gene_222 and gene_77328, all genes had their clear functions in photo response and photosynthetic processes. Interestingly, the targets of gene_77328 enriched in WD40 domaincontaining genes, involved in multi-cellular processes (Fig 8). Based on the metabolism pathway analysis of T2/T3 DEGs (FigS 4), the differentially altered metabolic pathways mainly focused on the lipid, carbohydrate, amino acid and energy dynamics.

**Fig 6.**
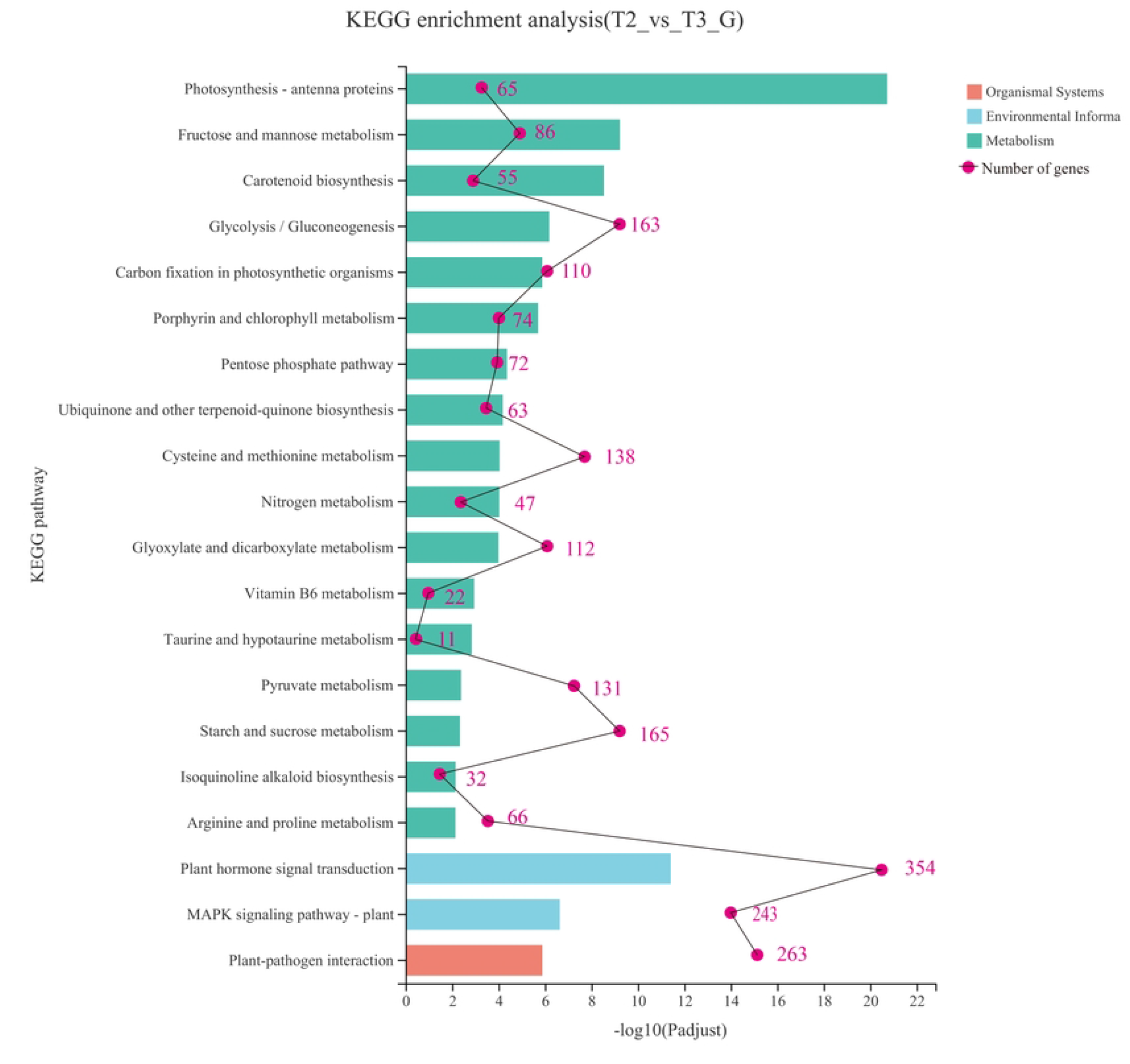
KEGG pathway enrichment analysis. Pathway enrichment of DEGs between T2 and T3 leaves. The number of DEGs was indicated by the purple Arabic numerals.

**Fig 7.**
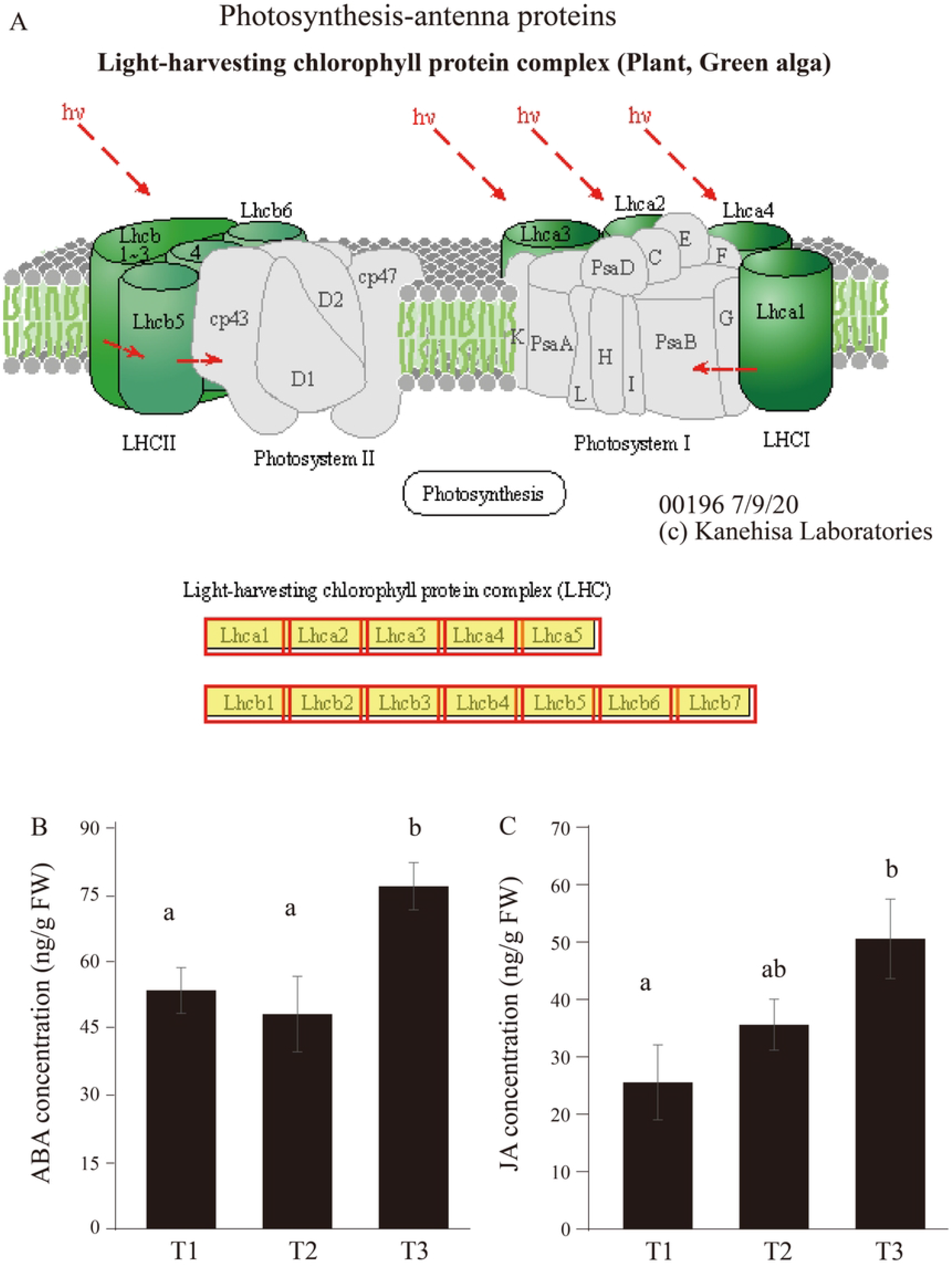
Differentially expressed genes encoding “photosynthesis-antenna proteins” and the hormone dynamics in T3 leaves. A, The red box indicates the upregulated DEGs. The schematic diagram of photosynthesis protein complex was cited from the map00196 in the KEGG database (https://www.genome.jp/pathway/map00196). ABA (B) and JA (C) concentrations in CX-26 cigar leaves. n=4. The significant differences were analyzed with one-way ANOVA.

**Fig 8.**
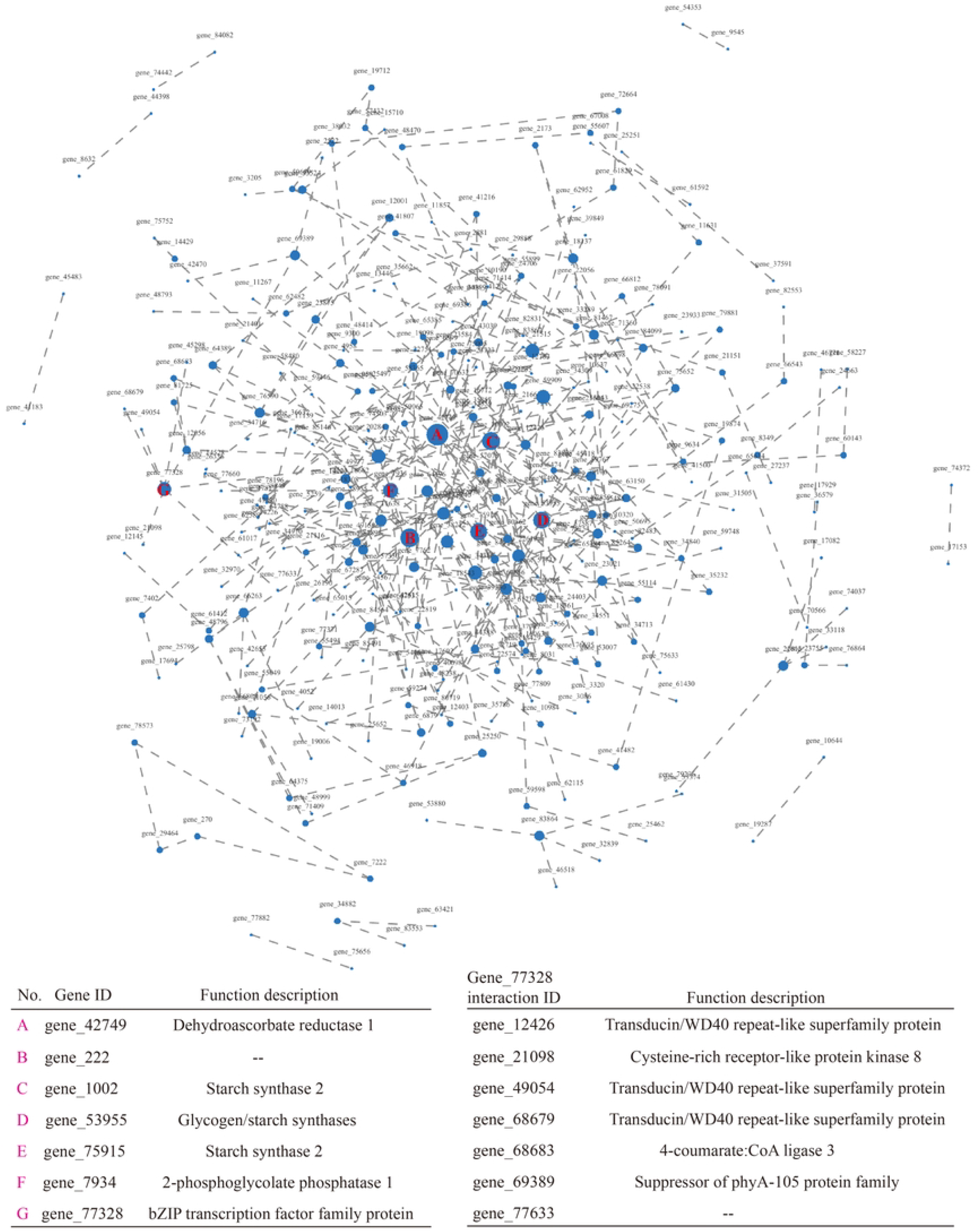
Gene regulatory network in the T3 leaves. Blue filled circles indicate the genes which were linked by dash lines, respectively. The larger of circle size, the stronger of importance. The top genes were listed in the left table using purple A->G letters and the interaction genes of gene_77328 were also listed in the right table.

## Discussion

Leaf senescence is a readout of diverse internal and environmental signals and occurs usually in the mature leaf, or occasionally exists in the young leaf under specific biotic and/or abiotic stress. This readout involves the highly dynamic processes at multiple regulatory layers from nucleic acid to protein (transcription, post-transcription, translation and post-translation and so on) and the metabolic alteration (phytohormones, amino acids, carbohydrates and so on).

In CX-26 cigar tobacco leaf, the visible color changes indicate the dramatic metabolic alterations in physiological level, marking a post-mature growth state. The significant reduction of SPAD value and the moisture content in T3 leaves (Fig 1) might be the beginning of senescence-associated chlorophyll degradation. This result suggests that the 71-71-day growth after transplanting is a key time point that triggers a growth transition from mature to post-mature of CX-26 middle leaves. In agricultural production practice, the maturity of cigar tobacco leaves adjudgment mainly depends on the experiences of the growers. Thus, the visible color changes of cigar leaves represent the late phase of maturity. The transcriptomic analysis is a helpful tool, providing a massive understanding of the gene transcriptional dynamics in cells. For example, important regulatory networks involved in the development dynamics of tobacco leaves were identified through transcriptomic analysis [7]. We performed the transcriptomic analysis to explore the early gene regulatory events involved in this growth state transition. About 80% of genes were detected in a genome scale in CX-26 middle leaves (Fig 2-3). Consistent with the reduction of SPAD value in T3 leaves, the DEGs of T1 and T2 leaves were fewer and PCA analysis showed an overlapped pattern of T1 and T2 expression, while T3 leaves had massive DEGs from T1 and T2 leaves and unique PCA pattern (Fig 4). These results indicate the distinct gene-regulatory networks in T3 leaves.

KEGG enrichment analysis point to the importance of photosynthesis (PS) antenna proteins, hormone signal transduction, MAPK signaling pathway and plant-pathogen interaction (Fig 6). Oxygenic photosynthesis is responsible for the great biomass of cigar tobacco leaves. The PS antenna proteins function in light capturing and energy transferring by binding Chla, Chlb, and carotenoids in the light-harvesting chlorophyll protein complexes embedded in the thylakoid membrane in green plants [13]. At the same time, the gene expression between T2 and T3 leaves involved in carotenoid biosynthesis, porphyrin and chlorophyll metabolism and carbon fixation in photosynthetic organisms was altered (Fig 6). These results are consistent with the GO annotations between T2 and T3 leaves such as the binding process, membrane component and metabolic process (Fig 5). Interestingly, either the LHC genes (Fig 7A) and the most of photosynthetic-related genes were upregulated (Fig 5B) supporting that T3 leaves would have stronger photosynthesis than the T2 leaves. However, the reduction of SPAD in T3 leaves can not provide sufficient chlorophyll content for photosynthesis. It has been well documented that ABA acts as a stress hormone regulating stomatal in plant response to drought stress to limit photosynthesis [14-16]. Recently, increasing attention had been paid to the ABA-mediated chlorophyll degradation and leaf senescence by transcriptional activation of chlorophyll catabolic genes and senescence-associated genes in many species [11, 17-20]. In this study, the ABA level was higher in T3 leaves than that of T1 and T2, suggesting that ABA might play a physiological role in the photosynthetic process involving chlorophyll degradation. It had been reported that MAPK6 plays a regulatory role in JA-mediated leaf senescence and suppressing the activity of MAPK6 delays leaf senescence [21-22]. ABA can activate MAPKs to mediate ABA signaling in plant cells such as reduced MAPK3 expression disrupts the ABA inhibition of stomatal opening [23]. This raises the key question of how MAPKs mediate the ABA-induced growth state changes of CX-26 leaves. Nevertheless, the elevated ABA, JA levels and MAPK signaling combined with the reduction of SPAD value suggest that chlorophyll degradation early occurs before other components decline such as starch synthesis, an important regulatory hub in the gene network (Fig 8). Dehdroascorbate reductase 1 gene (DHAR1) is the most important regulatory hub (Fig 8). In Arabidopsis, DHAR1 was induced by high light and the function reduction significantly impairs the modulation of cellular redox states under photooxidative stress [24]. The gene_77328 encodes a bZIP transcription factor protein, however, it did not directly starch synthase genes and LHC genes but some WD40 domain-containing family protein (Fig 8). WD40 domain has been widely found in eukaryotic proteins and acting as a scaffolding molecule for various protein activities. For example, the GIGANTUS1, a WD40 protein controls seed germination, growth and biomass in Arabidopsis [25]. The gene_77633 is the target of gene_77328 encoding the suppressor of phyA-105 protein (SPA) (Fig 8). It has been reported that SPA1 is a PHYA signaling intermediate, regulating PHYA signaling pathway involved in light-dependent photomorphogenesis, circadian rhythms and flowering time in plants [26-29]. This result suggests that gene_77328 might also play a critical role in the photosynthetic process. Another question needed to be considered is the “plant-pathogen pathway” enriched in T3 leaves. However, few literature discussed this issue on the maturity of post-mature tobacco leaves, though the ABA and JA would be the important players. It may chlorophyll degradation induced defense response overlaps the plant-pathogen and the hormones are the consensus regulator.

## Conclusions

In this study, we first found that the 71-75 days is important for the growth state transition accompanied by an evident SPAD reduction. Consistent with this, a transcriptome analysis of cigar leaves showed that a large number of genes had significant differential expression levels between 71-day leaves and 75-day leaves. The GO and KEEG analyses suggested that these differential expressing genes mainly enriched in the photosynthesis-related pathway and the hormone/MAPK signaling pathways, which was further supported by the gene regulatory network analysis and the elevated levels of ABA and JA in the 75-day leaves. Thus, we speculate that the ABA and JA dynamics in the different grown tobacco leaves might modulate the signal transduction pathways to affect the photosynthetic process, leading to a growth state transition. The detail of regulatory mechanisms can be further improved upon sustainable investigation.

## Materials and Methods

### Plant culture and experimental design

The Cigar (Nicotiana tabacum L.) Chuxue-26 (CX-26) was kept by our lab. The cigar seedlings were transplanted in Cuiba (E 109°47’42 “, N 30°27’27”), Enshi prefecture, Hubei province on May 16, 2021 and the apex was removed on July 2. Fertilizer application followed the description [30]. Middle leaves were sampled for the transcriptomic analysis on the July 22, 26 and 30, respectively.

### RNA extraction, library preparation and mRNA sequencing

Total RNA was extracted from leaves using TRIzol reagent (Invitrogen) without DNA contamination upon DNase I treatment (Takara). After quality control (OD26/280=1.8-2.2, OD260/230≥2.0, RIN≥6.5, 28S:18S≥1.0, >1 μg), RNA sample was used for library preparation. A total 1 μg RNA was used to construct the RNA-seq library using the TruSeq RNA sample preparation kit (San Diego, Ca). mRNA was isolated by oligo(dT) beads and then fragmented firstly. The cDNA was synthesized using a SuperScript double-stranded cDNA sysnthesis kit (Invitrogen, CA). The cDNA was subjected to end-repair, phosphorylation and ‘A’ base addition. A 15 PCR cycles was carried out for library size selection of target fragments and then the library was sequenced with the Illumina HiSeq xten/NovaSeq 6000 sequencer (2×150 bp read length).

### Read mapping, differential expression analysis and functional enrichment

The raw reads were trimmed and quality controlled on the web servers of SeqPrep (https://github.com/jstjohn/SeqPrep) and Sickle (https://github.com/najoshi/sickle). The clean reads were further aligned to reference genome with the HISAT2 software [31]. Then, the mapped reads were assembled by String Tie in a reference-based approach [32]. The value of each transcript expression was used to identify differential expression genes (DEGs) with the Q value ≤0.05, |log2FC|>1. The Goatools (https://github.com/tanghaibao/Goatools) and KOBAS (http://kobas.cbi.pku.edu.cn/home.do) were used to perform functional-enrichment analysis including GO terms and KEGG. These analysis tools were integrated in the Majorbio Cloud (Ren et al., 2022)[33].

### Moisture content and SPAD analysis

Cigar leaves harvested from field were immediately weighted biomass (M1), the dry weight (M2) was measured after 24 h of drying at 65 ºC. The moisture content was calculated due to the formula: 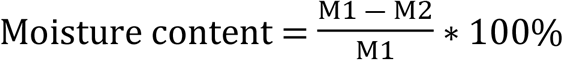. The SPAD value was recorded using the SAPD 502 meter.

### Hormone measurement

The method of hormone measurement followed the description by (Liu et al., 2012)[4]. Briefly, 0.1 g of fresh leaves was grinded well under liquid nitrogen condition and subsequently fully mixed with 750 μL pre-cooling buffer (methanol: H_2_O: acetic acid, 80: 19: 1, V/V/V) supplemented with internal standards, 10 ng ^2^H_6_ABA, 10 ng DHJA. The tubes were fixed on the shaker with 300 r/min shock at 4 ° C overnight. Tubes were centrifuged 10 min with 13000 rpm at 4 ° C and the supernatants were separated with a new tube. The precipitate was added 400 μL pre-cooling buffer without internal standards for another 4h of shaking at 4 ° C, and then the centrifuged supernatants were separated and merged with first supernatants. The supernatants were filtered with 0.22 μm membrane and dried with nitrogen gas. Then the tubes were added 200 μL 50% methanal to dissolve hormones for 3-6 h at 4 ° C. After 15 min of centrifuge with 13000 rpm at 4 ° C, 150-180 μL supernatants were isolated and stored in bottle for mass spectrometry. The hormone level analysis was carried out by the UFLC-ESI-MS.

### Statistical analysis

The significant differences among T1, T2 and T3 leaves were analyzed by the SPSS software with one-way ANOVA (P=0.05).

## Acknowledgments

We would like to thank Yu Luo (Huazhong Agricultural University) for her technical assistance.

## Supporting information

**S1 Fig. Differentially expressed genes involved in “plant hormone signal transduction” in T3 leaves**. The red and blue boxes indicate the upregulated and downregulated DEGs, respectively. yellow and green rectangles indicate the annotated and new genes, respectively. The schematic diagram of photosynthesis protein complex was modified from the map04075 in the KEGG database (https://www.genome.jp/pathway/map04075).

**S2 Fig. Differentially expressed genes involved in “MAPK signaling pathway-plant” in T3 leaves**. The red and blue boxes indicate the upregulated and downregulated DEGs, respectively. yellow and green rectangles indicate the annotated and new genes, respectively. The schematic diagram of photosynthesis protein complex was modified from the map04016 in the KEGG database (https://www.genome.jp/pathway/map04016).

**S3 Fig. Differentially expressed genes involved in “pathogen interaction” in T3 leaves**. The red and blue boxes indicate the upregulated and downregulated DEGs, respectively. yellow and green rectangles indicate the annotated and new genes, respectively. The schematic diagram of photosynthesis protein complex was modified from the map04626 in the KEGG database (https://www.genome.jp/pathway/map04626).

**S4 Fig. Metabolic pathway. The picture represents pathways annotated by gene sets of T2 and T3 leaves**. The red lines indicate the metabolic pathways that differentially altered between T2 and T3. Line wide represents the fold value of DEGs.

